# Fusion of blood vessel organoids with human pancreatic islets improves insulin response over time

**DOI:** 10.1101/2024.01.15.575704

**Authors:** Emily Tubbs, Mahira Mehanovic, Mélanie Lopes, Clément Quintard, Stéphanie Combe, Mathieu Armanet, Thomas Domet, Julia Sabatier, Sabrina Granziera, Josef Penninger, Delphine Freida, Xavier Gidrol

## Abstract

Pancreatic islet transplantation is a promising treatment strategy for type 1 diabetes, however there are still major challenges to overcome, including vascularization. Novel strategies for the generation of prevascularized islets with native microvessels have reported improved islet functionality, vascularization and engraftment emphasizing integral role of microvascular bed. Recently a new model of self-organizing three-dimensional human blood vessel organoids (BVOs) has been developed from human pluripotent stem cells (hPSCs), composed of both endothelial and mural cells. BVO recapitulate key features of human microvasculature such as formation of vascular network, vascular lumen and basement membrane, and have been shown to be perfusable. Here, we report a new strategy to construct prevascularized islets by fusion with hPSC-derived BVOs. We demonstrate that islets and BVOs in co-culture leads to fusion and improved insulin secretion over time, on two independent human islet donors, suggesting a new therapeutic approach for pancreatic islet transplantation and type 1 diabetes modeling.

## Introduction

Pancreatic islet transplantation is a promising treatment strategy for type 1 diabetes, however there are still major challenges to overcome, including vascularization. Physiologically, pancreatic islets have rich microvascular network and high metabolic rate, receiving fivefold higher blood flow than the exocrine pancreas^1^. However, microvascular network consisting of arterioles, capillaries, and venules is dissociated during the deceased donor islet isolation, contributing to poor engraftment outcomes with 25-50% of islet survival post-transplant^2^. A prompt and adequate blood supply during immediate post-transplant period is obstructed since graft revascularization process can take several days to weeks^3^. During this period, avascular islets depend on oxygen and nutrients supply by diffusion leading to central islet necrosis induced by ischemia upon intrahepatic islet transplantation^4^. Different strategies have emerged in a quest for improving islet survival including alternative sites of implantation and more recently bioengineering prevascularized islets with endothelial and proangiogenic supporting cells^5,6^. Nevertheless, the later approach often utilizes islets including primary endothelial cells from human umbilical cord lacking mural cells that are needed for proper blood vessel function. Novel strategies for the generation of prevascularized islets with native microvessels have reported improved islet functionality, vascularization and engraftment emphasizing integral role of microvascular bed^7,8^. Recently a new model of self-organizing three-dimensional human blood vessel organoids (BVOs) has been developed from human pluripotent stem cells (hPSCs)^9^. Composed of both endothelial and mural cells, BVO recapitulate key features of human microvasculature such as formation of vascular network, vascular lumen and basement membrane. Here, we report a new strategy to construct prevascularized islets by fusion with hPSC-derived BVOs. We demonstrate that islets and blood vessel organoids in co-culture leads to fusion and improved insulin secretion over time, on two independent islet donors.

## Materials and methods

### Human islets

The human pancreatic islets were provided by the Laboratory of Cell Therapy for Diabetes (LTCD), PRIMS facility, Institute for Regenerative Medicine and Biotherapy (IRMB), University hospital of Montpellier, Montpellier, France (authorization no.:AC-2019-3555); as well as the Cell Therapy Unit, AP-HP, Saint Louis Hospital, Paris, France, (authorization no.:AC-2020-4101). Human islets were isolated from pancreas from two brain-dead donors for therapeutic purposes. The procedure was approved by the National Agency for Medicines and Health Products Safety in accordance with the French bioethics law (Act No.2004-800;Act No.2011-814). After confirmation by the Agence de la Biomédecine, islets not suitable for transplantation are redirected to the cell therapy units’ scientific platform.

Human islet deceased donor information:

|  | Age | Sex | BMI (kg/m<sup>2</sup>) | Islet isolation center |
| --- | --- | --- | --- | --- |
| Donor 1 | 55 | M | 25.1 | LTCD Montpellier |
| Donor 2 | 52 | F | 25 | AP-HP Paris |

Islets were cultured in complete medium composed of CMRL 1066 medium, 1 g/L Glucose (Gibco, ThermoFisher Scientific, Waltham, Massachusetts, USA) supplemented with 10% FBS, 10 mM Hepes, 1 mM Sodium Pyruvate, 2 mM L-Glutamine, 100 mg.UI.mL^−1^ Penicillin, 100 μg mL^−1^ <u>streptomycin,</u> in a 5% CO_2_ humidified atmosphere at 37 °C.

### Blood vessel organoids (BVOs)

3D human blood vessel organoids were generated from human induced pluripotent stem cells (hiPSCs) as previously described^9^. In brief, stem cell colonies were dissociated using Accutase (Sigma-Aldrich, St. Louis, Missouri, United States) to get a single cell suspension. 600000 single stem cells were seeded per well (500 cells/microwell) of AggreWell™400 (STEMCELL Technologies, Vancouver, Canada) plates in aggregation media with 50 μM Y27632 (STEMCELL Technologies). Mesoderm induction and sprouting was induced directly in the AggreWell™400 plates by carefully changing the media with a p1000 pipette making sure to not disturb the cell aggregates in the microwells. For Collagen I-Matrigel embedding, organoids were collected from the AggreWell™400 plate by vigorously pipetting up and down with a cut p1000 tip close to the bottom of the well. Collected organoids were embedded in a 12-well plate (approx. 100 organoids/12-well), and subsequently cut out and singled into low attachment 96 U-well plates 4-5 days after embedding as previously described^9^. No alterations were made to any of the BVO differentiation media. The BVOs were maintained in a differentiation medium containing 15% FBS (Gibco, ThermoFisher Scientific), 100 ng/ml VEGF-A and 100 ng/ml fibroblast growth factor 2.

### Co-culture of human islets and BVOs

Native human islets were individually placed into a U-shaped 96-well microplate with an ultra-low attachment surface (Corning, New York, USA), and one single BVO was added per well. Islets and BVOs were cultured in complete islet medium mixed with the differentiation BVO medium (ratio 1:1). On the day of the co-culture, control islets medium was also changed with this same medium mix (ratio 1:1) to circumvent the sole effect of medium.

### Immunofluorescence staining

For immunofluorescent staining, the cells were fixed by Image-iT Fixative Solutions (ThermoFisher Scientific, R377814) for 30min to 2h at room temperature at 600 rpm, rinsed with PBS and stored at 4°C. The cells were permeabilized by a blocking buffer containing FBS (ScienCell Research Laboratories, Carlsbad, California, USA, 0010), sodium deoxycholate solution, Triton X-100 (Sigma-Aldrich, T8787), Tween 20 (Sigma-Aldrich, P1379) and Bovine Serum Albumin (Sigma-Aldrich, A9647) overnight at RT, 600 rpm. Primary antibodies rabbit anti-human CD31 (Abcam, Cambridge, United Kingdom, ab134168) and mouse anti-human glucagon (Abcam, ab10988) were diluted at 1:1000 in blocking buffer incubated 48h at 4°C, 600 rpm. After 3 washes in PBST (0.05% Tween), FITC goat anti-rabbit (Inerchim, Montluçon, France, 111-095-144) and goat anti-mouse Cy3-conjugated secondary antibodies (Interchim, 115-165-003) were diluted at 1:1000 in blocking buffer and incubated for 48h at 4°C, 600 rpm. After 3 washes in PBST, nuclear counter-staining using Hoechst 33342 (ThermoFisher Scientific, H-1399) was carried out according to a routine protocol for 2h at room temperature.

### Acquisition with light sheet fluorescence microscopy

After immunofluorescence staining, the cells were embedded into a cylinder of agarose gel then transparentized using a solution of CUBIC R2 overnight, and visualized by Light sheet fluorescence microscopy (LSFM, Z1 ZEISS); detection = EC Plan-Neofluar 5x/0.16 foc (WD=10.5 mm); illumination =2 LSFM 5x/0.1 foc objectives; space between different Z stacks=1.869-4.110 μm; pixel size= 0.37-0.91 μm; Light sheet thickness= 3.91-9.90 μm; using ZEN 3.1 software for Light sheet). 3D reconstructions were done on Arivis vision 4D software (v2.12.6x64).

### Glucose stimulated insulin secretion (GSIS) assays

Native human islets, or co-cultures with BVO individually placed in a 96-well plate with ultra-low attachment surface (Corning) were incubated at 37°C, 95% O2, 5% CO2 for 1h in 60 μL KREBS buffer 1% BSA, 2.8 mM glucose (Sigma-Aldrich, 49163) as pre-incubation. Thereafter, cells were incubated for 1h with 2.8 mM glucose solution (low glucose solution) and for 1h with 16.7 mM glucose solution (high glucose solution). After each incubation step, supernatants were collected and stored at −80 °C. Insulin concentration in each collected supernatant was measured in duplicate using STELLUX Chemiluminescence Human Insulin ELISA (ALPCO, Salem, New Hampshire, United States). A stimulation index (SI) was obtained by calculating the ratio of insulin measured between high and low glucose stimulation [high glucose solution]/[low glucose solution].

### Statistical analysis

Results are expressed as mean ± SEM, and statistical significance was defined as a value of *\*P*<0.05, *\*\*P*<0.01, *\*\*\*P*<0.001. Student t test was used for statistical analysis using GraphPad Prism 9 (GraphPad Software Inc., San Diego, CA, USA).

## Results

### Human islets and blood vessel organoids fuse together

Human islets were co-cultured with BVOs, with one individual islet paired with one individual BVO (Figure 1). Initially, BVOs were generated from hiPSCs in 3D collagen I-Matrigel matrix and collected for co-culture with human islets, using a medium mix of each cell types. The co-culture kinetics was monitored daily by bright-field microscopy, and engulfment of the human islet and BVO was observed at day 3. At day 5, complete physical fusion was observed between islet and BVO (Figure 1B) in each well of co-culture (n=5-8), with 100% fusion success. This was confirmed with 2 different human islet donors.

**Figure 1:**
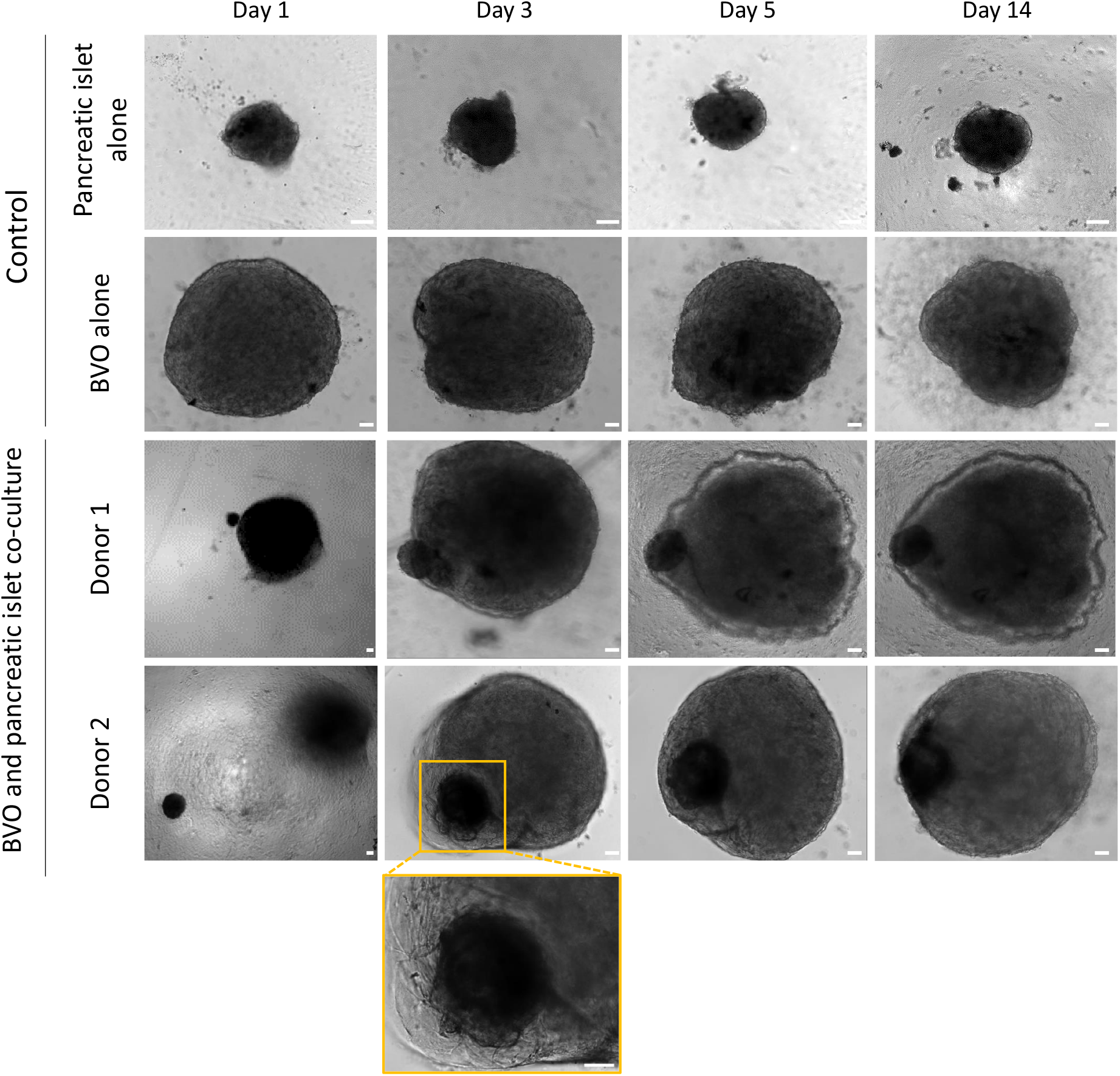
Auto-organization and fusion of human islet and blood vessel organoid. On day 1, a human islet and a blood vessel organoid (BVO) are put in co-cultured. On day 3 partial engulfment is observed. On day 5, the human islet is fully embraced by the BVO, and remains fused until day 14. Images obtained by brightfield microscopy. Scale bar = 100μm.

To understand the sequence of events during the fusion, an in-depth analysis and 3D observation of whole samples were performed with light sheet fluorescence microscopy (LSFM) using islet specific marker glucagon and endothelial cell marker CD31 at day 3 and day 7 of co-culture. Maximum intensity projection of LSFM Z-stacks was used for 3D visualization of BVO and islet structures that were distinguished with CD31 and glucagon labeling, respectively (Figure 2A). Representative Z-stack images of CD31 and glucagon immunostaining were used to detect islet engulfment within the BVO at day 3 and at day 7 of co-culture, corresponding to complete fusion (Figure 2B). Single LSFM Z-stacks were used for detection of double positive regions, indicating islet invasion and localization within BVO. The 3D reconstruction of glucagon positive islets and CD31 positive BVOs was also visualized in a video (Video 1S). Taken together, these preliminary results on two different islet donors seem to confirm complete fusion of islets with BVOs, as well as invasion of the islet with endothelial cells.

**Figure 2:**
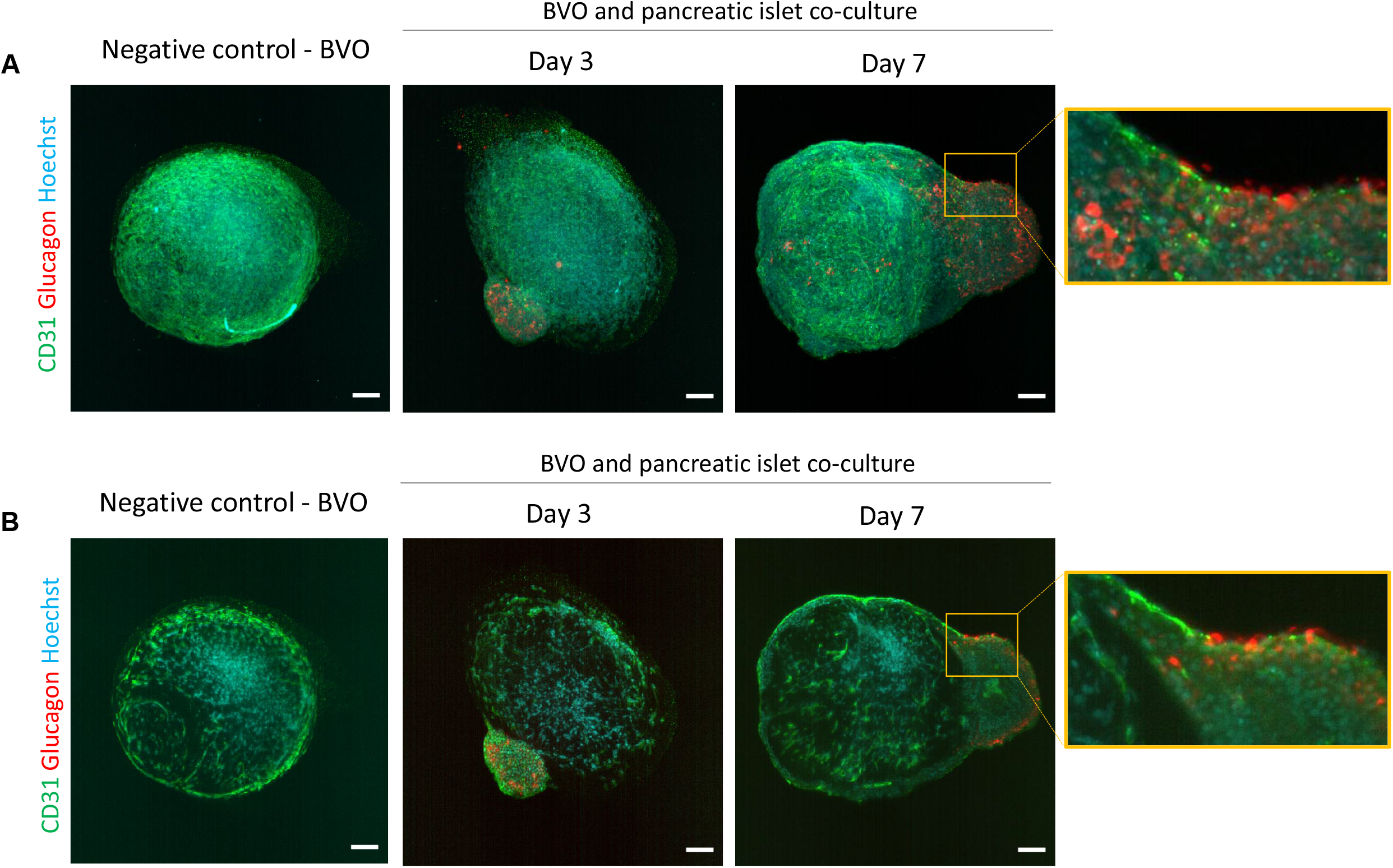
Fusion of human islet and blood vessel organoid. **A)** Maximum intensity projection of Z-stacks showing immunofluorescence labeling with CD31 (green) and glucagon (red) of BVO alone or with islet on day 3 and day 7 of co-culture. **B)** Single Z-stack section showing CD31 (green) and glucagon (red) labeling of BVO alone or with islet on day 3 and day 7 of co-culture. The islet and BVO fusion and penetration on day 7 is shown in orange boxes. Scale bar = 100μm.

### Co-culture with blood vessel organoids improve islet viability and functionality

Following the fusion of islets and BVOs, we first performed live and dead cell labeling with SYTO13 and propidium iodide (PI), showing that the co-culture conditions as well as the mix of medium used had no negative impact on viability at day 3 and day 7 (*data not shown*). Then we investigated the functionality of islets fused with BVOs by performing glucose stimulated insulin secretion assay at day 3 or day 5, and at day 7 and 14. We observed similar insulin secretion profiles for the two pancreatic islet donors.

Islets co-cultured with BVOs had similar profiles compared to islets cultured alone, showing that BVOs had no negative impact on islets functionality (Figure 3A). In fact, we observed slightly higher insulin secretions levels when islets were co-cultured with BVOs for donor 1, as soon as day 3. Moreover, for donor 2 we observed significantly higher insulin secretion when exposed to high glucose concentration at day 7 of co-culture. These preliminary results suggest that co-culturing islets and BVOs could present a benefit on human islet functionality. To go further, the stimulation index was calculated as the ratio between insulin secretion after high and low glucose stimulations (Figure 3B), and as expected, islets from both donors showed a gradual decline in glucose responsiveness over the 14 days period studied when cultured alone. Strikingly, the glucose responsiveness was maintained and persisted longer in islets co-cultured with BVOs, with significantly higher SI on day 14 of co-culture.

**Figure 3:**
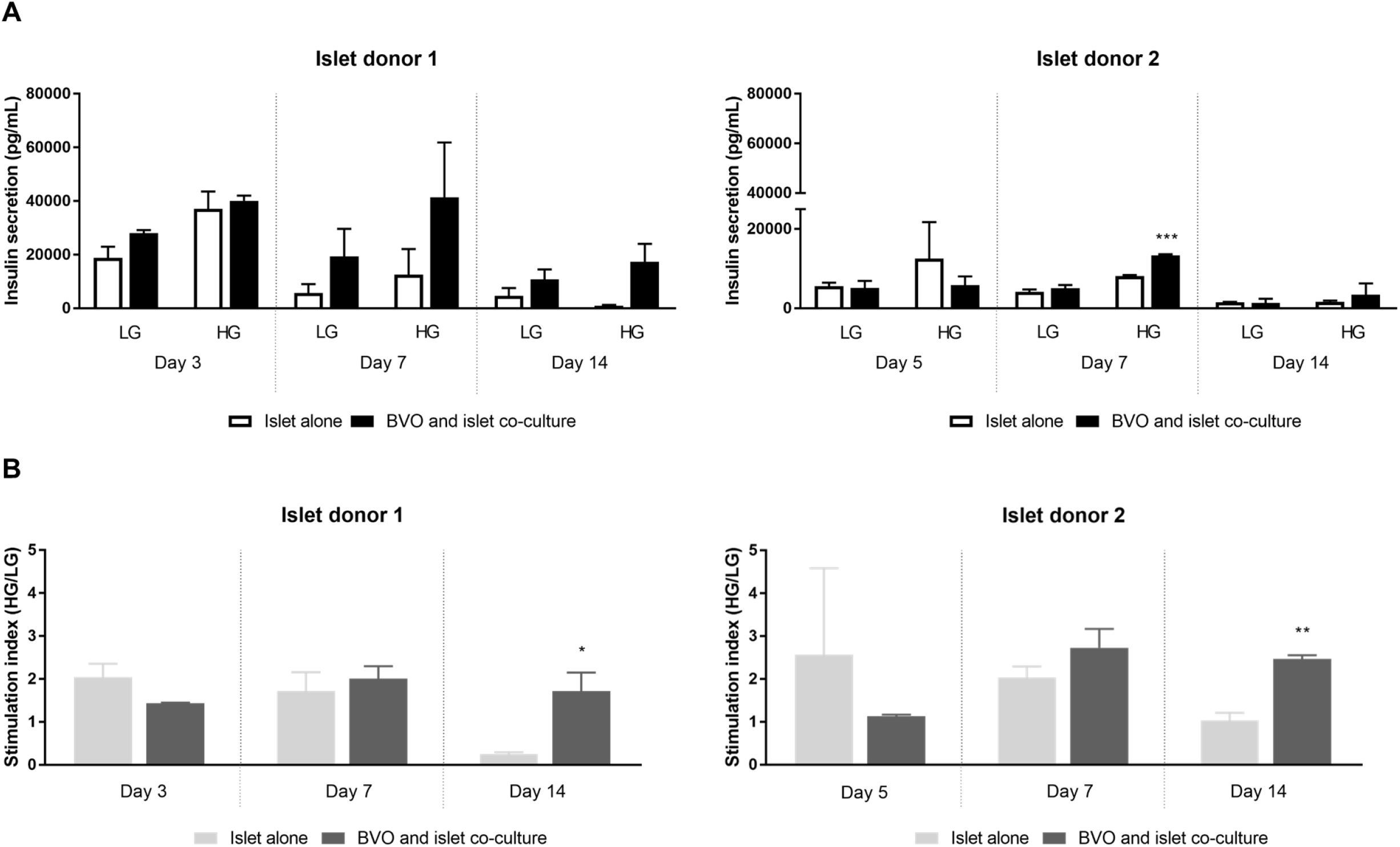
Improved functionality of human islet co-cultured with blood vessel organoid. Comparison of the insulin secretions of human islets co-cultured with blood vessel organoid (BVO) at day 3 or day 5; day 7 and 14 in response to low glucose solution (LG; 2.8mM) and then to high glucose solution (HG; 16.7mM), on 2 separate experiments corresponding to two independent human islet donors, each condition performed in triplicate **(A)** and relative stimulation index = [high glucose solution]/[low glucose solution] **(B)**.

Overall, our preliminary results show that the co-culture with BVOs maintains the insulin secretory function of the islet until day 14.

## Discussion

In this study, we present an islet prevascularization strategy wherein the co-culture of islet and BVO results in their fusion, with maintained islet functionality over-time. The fusion with BVO prevented the pancreatic islet from a gradual decline in insulin secretion classically observed when islets are cultured alone. To our knowledge this is the first proof-of-concept study of islet and BVO fusion.

Previous studies reported successful fusion of hPSC-derived brain and blood vessel organoids providing the brain organoid with robust vascular network-like structures simulating the *in vivo* developmental processes of the brain for further applications in various neurological disease^10–13^. This evidence suggests that BVOs represent a functional microvasculature model that can be used in multiple organoid fusion settings.

Our study demonstrates the utilization of BVOs for islet prevascluarization, a strategy that could potentially improve islet transplantation outcomes. Interestingly, previous studies have shown that co-transplantation of human islets with microvascular fragments (MVFs) isolated from adipose tissue significantly improve islet survival and function post-transplantation^8^. The MVFs have been shown to be a justified source for islet vascularization as they provide functional fragments of arterioles, capillaries, and venules. Islets secrete VEGF-A to activate endothelial cells and vascularization, while laminin-rich vascular basement regulates insulin gene expression^14^. These interactions between the microvasculature and islets are crucial for islet vascularization and function. The study by Nalbach et al. (2021) showed that fusion of MVFs with human islets improves their vascularization, viability and functionality post-transplant^7^. However, the islet prevascularization with MVFs included human islet dissociation and re-aggregation, a common approach of prevascularization strategies which can often be detrimental to the islet physiology as well as labor-intensive and time-consuming^15^. In addition, the choice to re-aggregate islets with endothelial and stromal cells instead of microvasculature units can lead to their separation and migration out of islet aggregates^16,17^. During the first days of co-culture we observed fusion in all tested wells showing remarkable fusion efficiency. Moreover, we did not observe any islet escape from BVOs during 2 weeks of co-culture. Our vascularization strategy does not require pseudoislet formation, nor does it utilize microvascular fragments but instead employs complete microvasculature units.

A major limitation to the clinical application of fused BVO and islets so far, is the need for a collagen I-Matrigel matrix to generate BVOs. Interestingly, we cannot exclude that the remaining collagen I-Matrigel matrix around the BVO could serve as an anchoring point for the islet before fusion, or influence vascularization. Other studies have shown a beneficial effect of collagen hydrogels and MVFs on islet vascularization and function^18^. Other matrixes could certainly be used, but they require further investigations for applications in regenerative medicine. Clinical co-transplantation of deceased donor islets fused with BVOs would be possible with simple co-culture to allow their fusion in a few days.

Recent advancements in stem cell-derived pancreatic islets are an extremely encouraging source of insulin secreting cells overcoming human organ donor scarcity for islet transplantation^19^. Interestingly, the hiPSC-derived pancreatic islets and BVOs could be generated from same stem cell source, such as hypoimmunogenic stem cell lines to avoid rejection or from patient-derived hiPSCs. This opens a path for allogeneic or autologous co-transplantation of fused stem cell-derived islets and blood vessel organoids.

Interestingly, BVOs have also been shown to enable perfusion into an organ-on-chip model^20^, highlighting the tremendous possibilities of disease modeling using both the fusion and perfusion capacities of BVOs.

Taken together, the prevascularization of pancreatic islets seems to be a promising new approach to improve islet transplantation for patients, and hopefully the fusion strategy suggested in this paper between BVOs and islets will enable a swift and efficient approach applicable to islet transplantation for patients.

## Supporting information

Graphical abstract

## Abbreviations

BMI: body mass index
BVO: blood vessel organoid
CD31: cluster of differentiation 31
ELISA: enzyme-linked immunosorbent assay
FBS: fetal bovine serum
FITC: fluorescein isothiocyanate
GSIS: glucose stimulated insulin secretion
hiPSC: human induced pluripotent stem cells
hPSCs: human pluripotent stem cells
LSFM: light sheet florescent microscopy
MVF: microvascular fragment
PBST: Phosphate Buffered Saline with Tween
SI: stimulation index
VEGF: vascular endothelial growth factor

## Acknowledgements

This work was supported by the CEA “OOC inflexion” and “Focus organoids”. We thank the microscopy facility MuLife of IRIG/DBSCI, funded by CEA Nanobio and GRAL LabEX (ANR-10-LABX-49-01) financed within the University Grenoble Alpes graduate school CBH-EUR-GS (ANR-17-EURE-0003).

## Data availability statement

The data that support the findings of this study are available from the corresponding author upon reasonable request.

## References

1. Brusko TM, Russ HA, Stabler CL. Strategies for durable β cell replacement in type 1 diabetes. Science. 2021;373(6554):516–522. doi:10.1126/science.abh1657

2. Citro A, Moser PT, Dugnani E, et al. Biofabrication of a vascularized islet organ for type 1 diabetes. Biomaterials. 2019;199:40–51. doi:10.1016/j.biomaterials.2019.01.035

3. Brissova M, Powers AC. Revascularization of Transplanted Islets. Diabetes. 2008;57(9):2269–2271. doi:10.2337/db08-0814

4. Giuliani M, Moritz W, Bodmer E, et al. Central Necrosis in Isolated Hypoxic Human Pancreatic Islets: Evidence for Postisolation Ischemia. Cell Transplant. 2005;14(1):67–76. doi:10.3727/000000005783983287

5. Wassmer CH, Lebreton F, Bellofatto K, et al. Bio-Engineering of Pre-Vascularized Islet Organoids for the Treatment of Type 1 Diabetes. Transpl Int. 2022;0. doi:10.3389/ti.2021.10214

6. Takahashi Y, Sekine K, Kin T, Takebe T, Taniguchi H. Self-Condensation Culture Enables Vascularization of Tissue Fragments for Efficient Therapeutic Transplantation. Cell Reports. 2018;23(6):1620–1629. doi:10.1016/j.celrep.2018.03.123

7. Nalbach L, Roma LP, Schmitt BM, et al. Improvement of islet transplantation by the fusion of islet cells with functional blood vessels. EMBO Mol Med. 2021;13(1):e12616. doi:10.15252/emmm.202012616

8. Aghazadeh Y, Poon F, Sarangi F, et al. Microvessels support engraftment and functionality of human islets and hESC-derived pancreatic progenitors in diabetes models. Cell Stem Cell. 2021;28(1):1936–1949.e8. doi:10.1016/j.stem.2021.08.001

9. Wimmer RA, Leopoldi A, Aichinger M, Kerjaschki D, Penninger JM. Generation of blood vessel organoids from human pluripotent stem cells. Nat Protoc. 2019;14(1):3082–3100. doi:10.1038/s41596-019-0213-z

10. Song L, Yuan X, Jones Z, et al. Assembly of Human Stem Cell-Derived Cortical Spheroids and Vascular Spheroids to Model 3-D Brain-like Tissues. Sci Rep. 2019;9(1):5977. doi:10.1038/s41598-019-42439-9

11. Kong D, Park KH, Kim DH, et al. Cortical-blood vessel assembloids exhibit Alzheimer’s disease phenotypes by activating glia after SARS-CoV-2 infection. Cell Death Discov. 2023;9(1):1–13. doi:10.1038/s41420-022-01288-8

12. Ahn Y, An JH, Yang HJ, et al. Human Blood Vessel Organoids Penetrate Human Cerebral Organoids and Form a Vessel-Like System. Cells. 2021;10(8):2036. doi:10.3390/cells10082036

13. Sun XY, Ju XC, Li Y, et al. Generation of vascularized brain organoids to study neurovascular interactions. Gleeson JG, Chin J, eds. eLife. 2022;11:e76707. doi:10.7554/eLife.76707

14. Nikolova G, Jabs N, Konstantinova I, et al. The Vascular Basement Membrane: A Niche for Insulin Gene Expression and β Cell Proliferation. Developmental Cell. 2006;10(3):397–405. doi:10.1016/j.devcel.2006.01.015

15. Wassmer CH, Bellofatto K, Perez L, et al. Engineering of Primary Pancreatic Islet Cell Spheroids for Three-dimensional Culture or Transplantation: A Methodological Comparative Study. Cell Transplant. 2020;29:0963689720937292. doi:10.1177/0963689720937292

16. Nguyen-Ngoc KV, Jun Y, Sai S, et al. Engineered Vasculature Induces Functional Maturation of Pluripotent Stem Cell-Derived Islet Organoids. Cell Biology; 2022. doi:10.1101/2022.10.28.513298

17. Jun Y, Kang AR, Lee JS, et al. Microchip-based engineering of super-pancreatic islets supported by adipose-derived stem cells. Biomaterials. 2014;35(7):4815–4826. doi:10.1016/j.biomaterials.2014.02.045

18. Salamone M, Rigogliuso S, Nicosia A, Campora S, Bruno CM, Ghersi G. 3D Collagen Hydrogel Promotes In Vitro Langerhans Islets Vascularization through ad-MVFs Angiogenic Activity. Biomedicines. 2021;9(7):739. doi:10.3390/biomedicines9070739

19. Hogrebe NJ, Ishahak M, Millman JR. Developments in stem cell-derived islet replacement therapy for treating type 1 diabetes. Cell Stem Cell. 2023;30(5):530–548. doi:10.1016/j.stem.2023.04.002

20. Quintard C, Jonsson G, Laporte C, et al. An Automated Microfluidic Platform Integrating Functional Vascularized Organoids-on-Chip. bioRxiv; 2021. doi:10.1101/2021.12.29.474327

