## Supplementary material for "Fusion of blood vessel organoids with human pancreatic islets improves insulin response over time": Graphical abstract

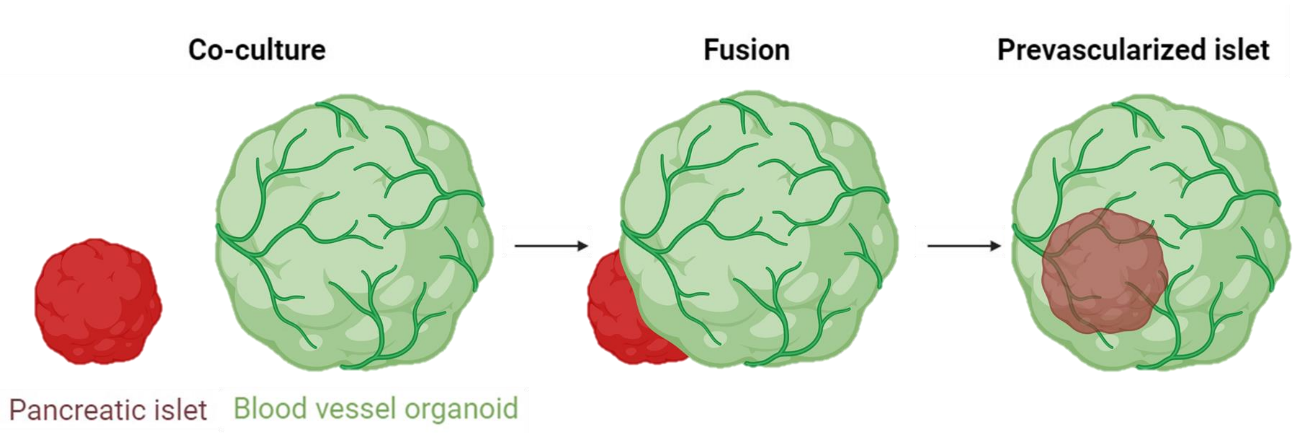


**Graphical abstract:** Schematic representation of islet and blood vessel organoid (BVO) co-culture leading to fusion.
