## Supplementary material for "Fusion of blood vessel organoids with human pancreatic islets improves insulin response over time": Supp data_video

### Slide 1
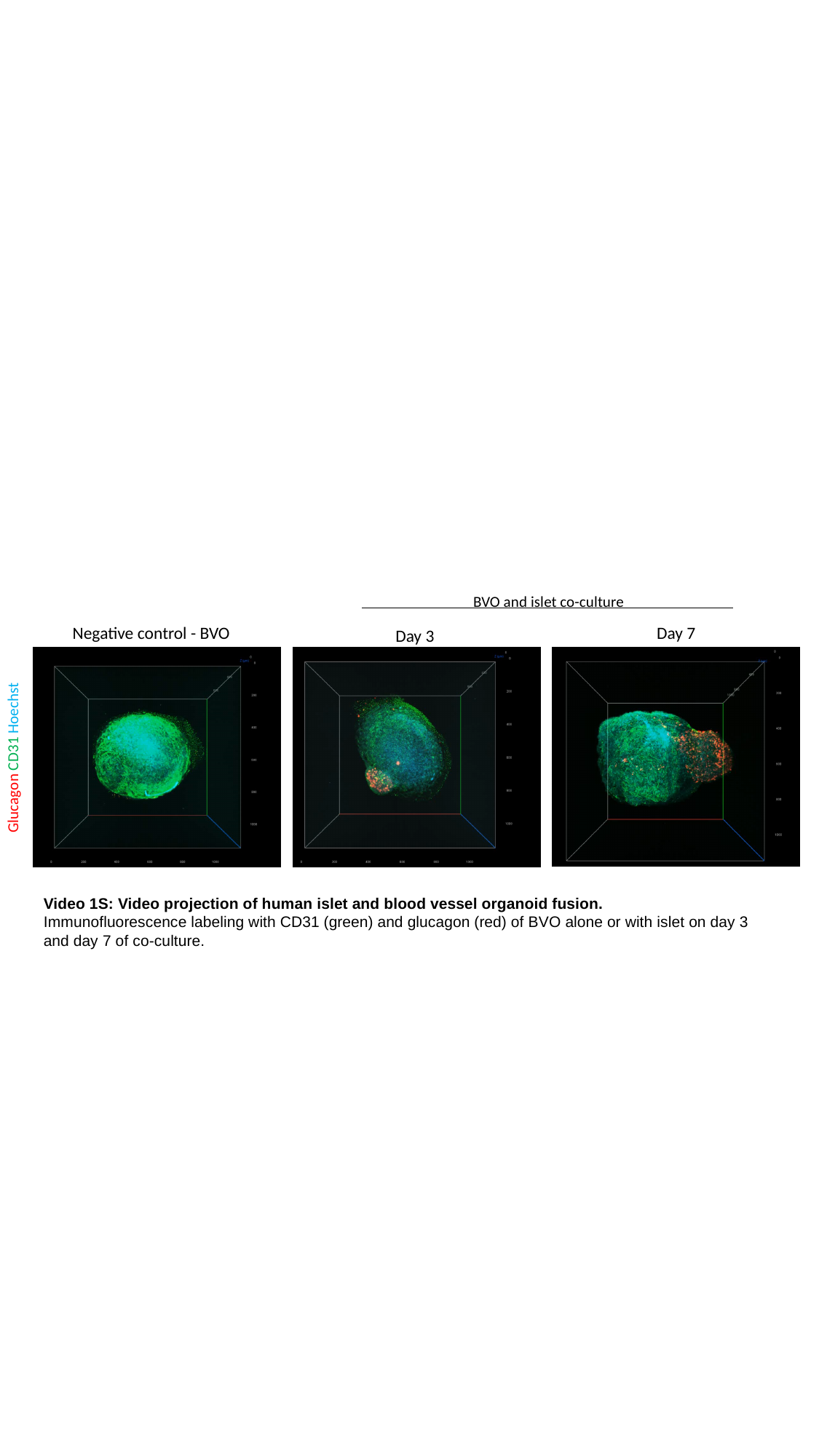

BVO and islet co-culture
Negative control - BVO
Day 7
Day 3
Glucagon CD31 Hoechst
Video 1S: Video projection of human islet and blood vessel organoid fusion.
Immunofluorescence labeling with CD31 (green) and glucagon (red) of BVO alone or with islet on day 3 and day 7 of co-culture.
